## Supplementary Material for "Controllability boosts neural and cognitive signatures of changes-of-mind in uncertain environments"

### Controllability reveals defining features of information seeking

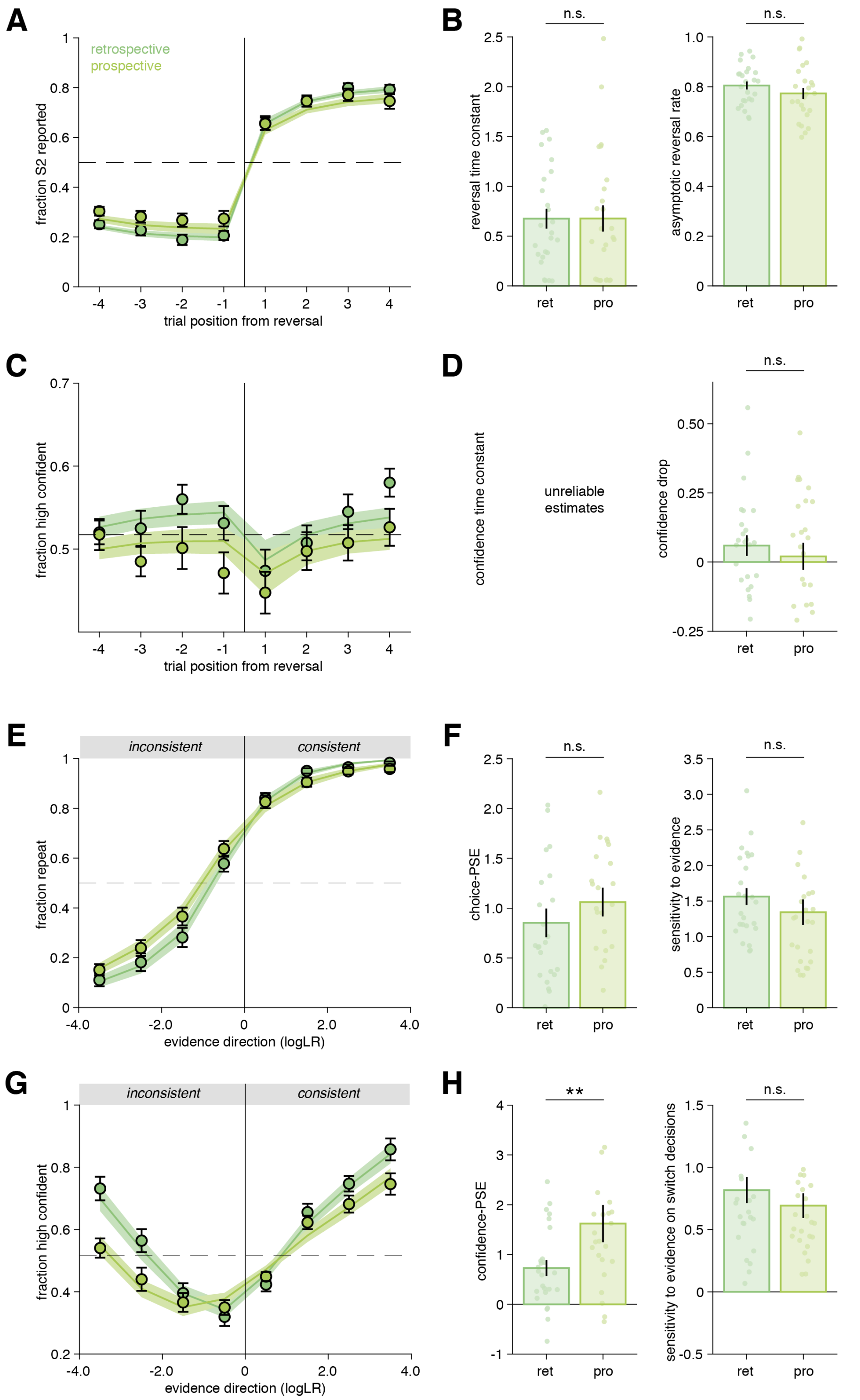


**Figure 2 supplement 1. Psychometric analysis of choice and confidence reversal and repetition curves in retrospective and prospective conditions (Experiment 3)**

**A**) Fraction of hidden state correctly reported as a function of trial number before and after a reversal. Vertical lines indicate the position of reversals. Horizontal dotted line indicates chance level. Circles and error bars indicate mean and SEM across participants (*N*=25). Shaded areas indicate mean and SEM of psychometric predictions from the best-fitting truncated exponential functions. **B**) Psychometric parameters ‘reversal time constant’ and ‘asymptotic reversal rate’ characterizing the fitted response reversal curve in A (see Methods). Circles indicate individual participants and bars and error bars indicate the mean and SEM across participants. n.s. not significant, paired t-tests. **C**)

Fraction of high confidence responses as a function of trial number before and after a reversal. Vertical lines indicate the position of reversals. Horizontal dotted lines indicate mean confidence. Circles indicate human data and error bars display S.E.M. across participants. Shaded areas indicate psychometric predictions from the best-fitting sigmoid functions. **D**) Psychometric parameter for the confidence reversal curve in C. Dots indicate individual participants. Because confidence drop is not different from zero in both conditions, confidence time constant are not reliably estimable and therefore are not presented. n.s. not significant, paired t-tests. **E**) Fraction of response repetitions as a function of whether the evidence was consistent (in favor of repeating) or inconsistent with the previous choice. Circles indicate human data and error bars display SEM across participants. Shaded areas indicate mean and SEM of psychometric predictions from the best-fitting truncated exponential functions. **F**) Psychometric parameters 'point of subjective equivalence' (choice-PSE) and 'sensitivity to evidence' (slope) characterizing the response repetition curves in E. n.s. not significant, paired t-tests. **G**) Fraction of high confidence responses as a function of whether the evidence was consistent or inconsistent with the previous choice. Circles indicate human data and error bars display within-subject S.E.M. Within-subject error bars are presented to allow a comparison between conditions without an influence of inter-individual variability about the use and calibration of the high and low confidence responses. Shaded areas indicate mean and S.E.M. of psychometric predictions from the best-fitting mixture of two sigmoid functions for repeat and switch trials respectively (see Methods). **H**) Psychometric parameters characterizing the confidence repetition curves in G. Left panel: 'confidence-PSE' (point of subjective equivalence) representing the quantity of evidence for which participants are as confident in their repeat decisions as in their switch decisions. **$p$<.01, Wilcoxon signed rank test (see Methods). Right panel: 'sensitivity to evidence' for the switch trials. Dots indicate individual participants and bars and error bars indicate the mean and SEM across participants. Panels **A**-**B**-**E**-**F** are reproduced from (Weiss et al., 2021).

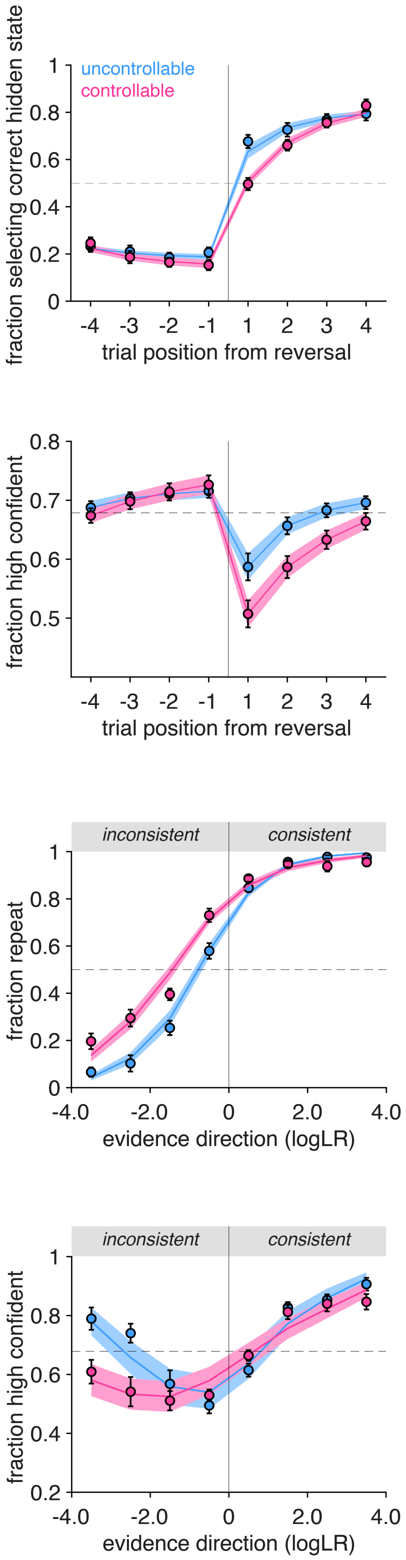


**Figure 2 supplement 2. Psychometric analysis of Experiment 2B**
**A**) Fraction of hidden state correctly reported as a function of trial number before and after a reversal. Vertical line indicates the position of reversals. Horizontal line indicates chance level. Shaded areas indicate mean and SEM of psychometric predictions from the best-fitting truncated exponential functions. **B**) Fraction of response repetitions as a function of whether the evidence was consistent (in favor of repeating) or inconsistent with the previous choice. Shaded areas indicate mean and SEM of psychometric predictions from the best-fitting sigmoid functions. **C**) Fraction of high confidence responses as a function of trial number before and after a reversal. Horizontal dotted lines indicate

mean confidence. Shaded areas indicate mean and SEM of psychometric predictions from the best-fitting sigmoid functions. In A, B and C, circles and error bars indicate mean and SEM across participants. **D**) Fraction of high confidence responses as a function of whether the evidence was consistent or inconsistent with the previous choice. Circles indicate human data and error bars display within-subject SEM Within-subject error bars allow a comparison between conditions without an influence of inter-individual variability about the use and calibration of high and low confidence responses. Shaded areas indicate mean and SEM of psychometric predictions from the best-fitting mixture of two sigmoid functions for repeat and switch trials respectively. Bars and error bars indicate mean and SEM across participants and dots indicate individual participants (*N*=18). Blue: uncontrollable (C-) condition, pink: controllable (C+) condition.

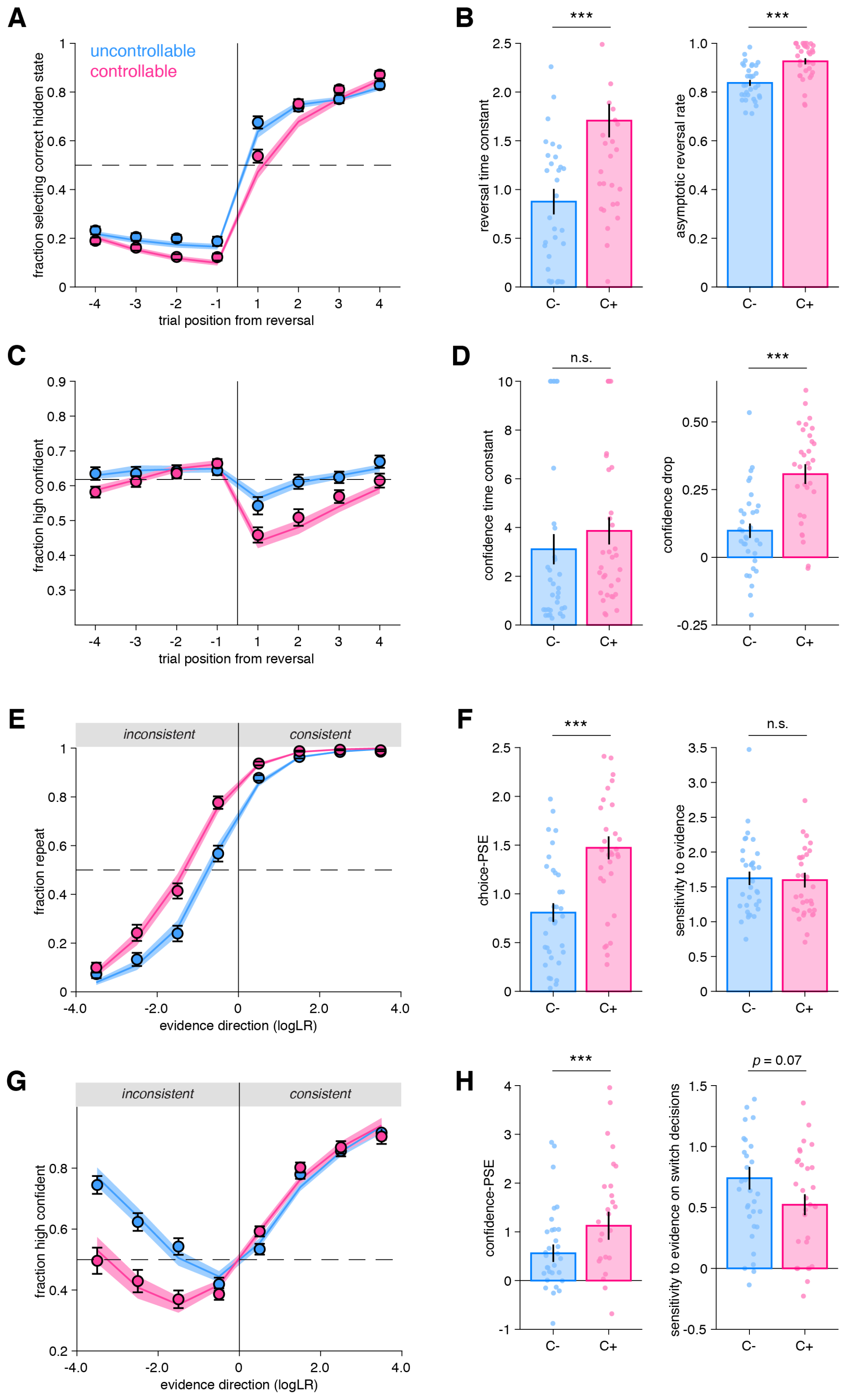


**Figure 2 supplement 3. Behavioral signatures of choice and confidence validate the computational model.** Simulations from the best-fitting parameters (shaded areas) in uncontrollable (C-, blue) and controllable (C+, pink) conditions (*N*=33, pooled across Experiments 1 and 2A). Same conventions as Fig. 2. **A**) Fraction of hidden state correctly reported as a function of trial number before and after a reversal. **B**) Psychometric parameters 'reversal time constant' (***$p$=9.5×$10^{-7}$, $t_{32}$=-

6.0) and ‘asymptotic reversal rate’ (***$p$=2.0×10$^{-7}$, $t_{32}$=-6.6) characterizing the fitted response reversal curve in A (paired $t$-tests). **C**) Fraction of high confidence responses as a function of trial number before and after a reversal. **D**) Psychometric parameters ‘confidence time constant’ and ‘confidence drop’ for the fitted confidence reversal curve in B. n.s. not significant, $p$=0.62, $t_{32}$=0.50; ***$p$=1.0×10$^{-5}$, $t_{32}$=-5.22; paired $t$-tests. **E**) Fraction of response repetitions as a function of whether the evidence was consistent or inconsistent with the previous choice. **F**) Psychometric parameters ‘point of subjective equivalence’ (choice-PSE) and ‘sensitivity to evidence’ (slope) characterizing the response repetition curves in E. ***$p$=1.7×10$^{-9}$, $t_{32}$=-8.3; n.s. not significant, $p$=0.43, $t_{32}$=0.79; paired $t$-tests. **G**) Fraction of high confidence responses as a function of whether the evidence was consistent or inconsistent with the previous choice. **H**) Psychometric parameters characterizing the confidence repetition curves in G. Left panel: ‘confidence-PSE’ (point of subjective equivalence) representing the quantity of evidence for which participants are as confident in their repeat decisions as in their switch decisions. ***$p$=.00033, $z$=-3.69, Wilcoxon signed rank test. Right panel: ‘sensitivity to evidence’ for the switch trials. $t_{32}$=1.86, $p$=.07, paired $t$-test.

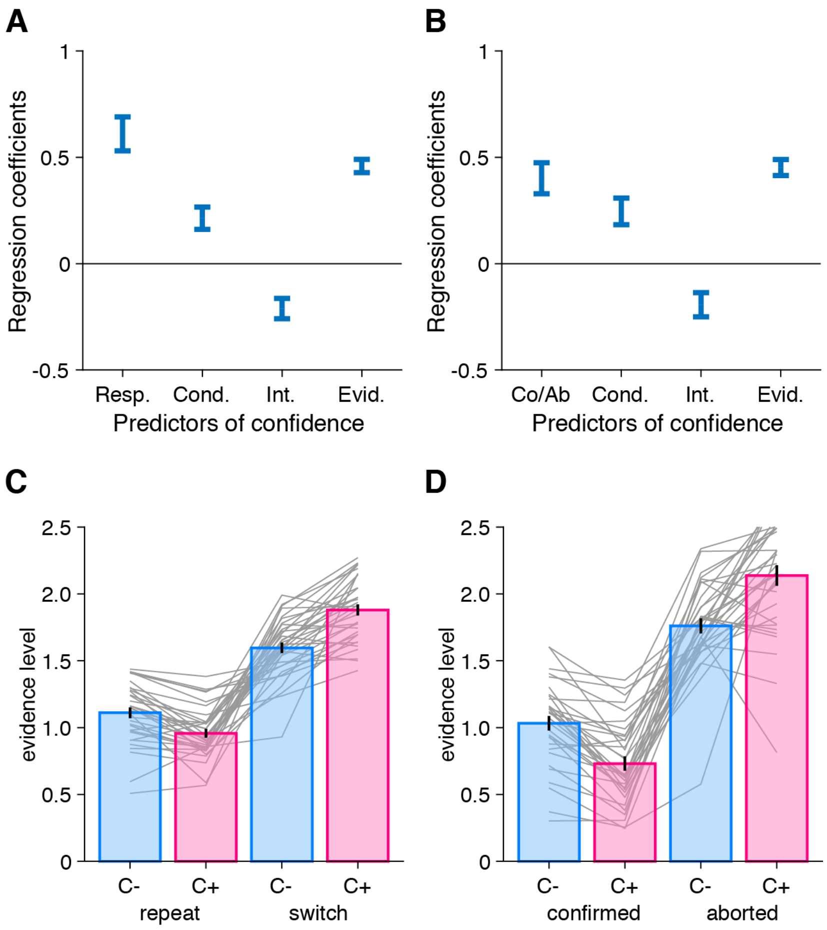


**Figure 3 supplement 1**. **A**: Logistic regression predicting confidence as a function of response type (Resp.) (repeat, switch), condition (Cond.) (C-, C+), their interaction (Int.), and the strength of evidence in favour of repeating the previous response (Evid.). **B**: Logistic regression predicting confidence as a function of whether the change-of-mind of the previous response was confirmed or aborted (Co/Ab), condition (Cond.) (C-, C+), their interaction (Int.), and the strength of evidence in favour of repeating the previous response (Evid.). **C**) Mean evidence level as a function of whether participants repeated their previous choice ("repeat") or changed their mind ("switch") in the uncontrollable (C-, blue) and controllable (C+, pink) conditions. **D**) Mean evidence levels following switch decisions that were confirmed or aborted i.e. when participants return back to their previous response in the two conditions. Note that individual variability is higher on the right panel because there are less events per participant in each of the cases. Bars and error bars indicate mean and SEM across participants and grey lines display individual data points (*N*=33).

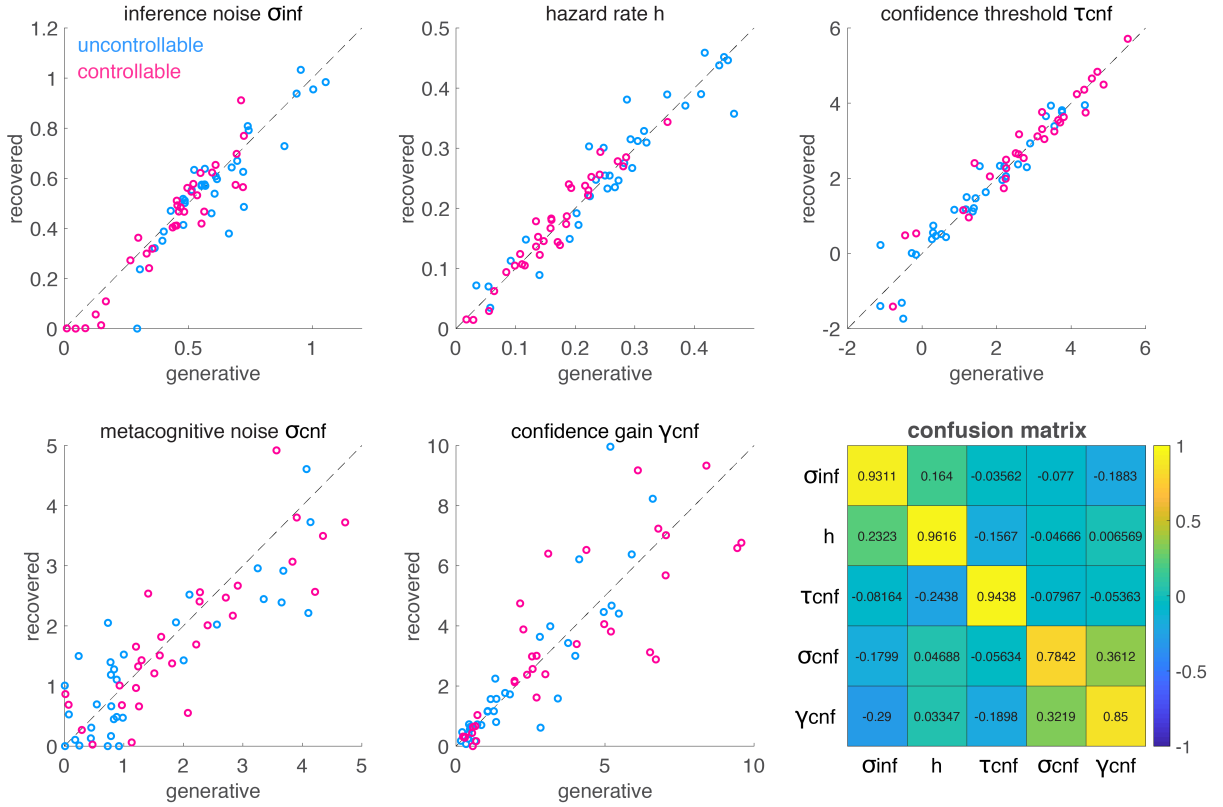


**Figure 4 supplement 1: Parameter recovery analysis.**
Pairwise correlations between generative and recovered inference noise, hazard rate, confidence threshold, metacognitive noise, and confidence gain parameters, in both conditions (blue: uncontrollable (C-); pink: controllable (C+)). The diagonal line corresponds to a very satisfactory recovery with strong correlations between generative and fitted parameters. Bottom right panel: Confusion matrix for the five parameters indicating a satisfactory recovery.

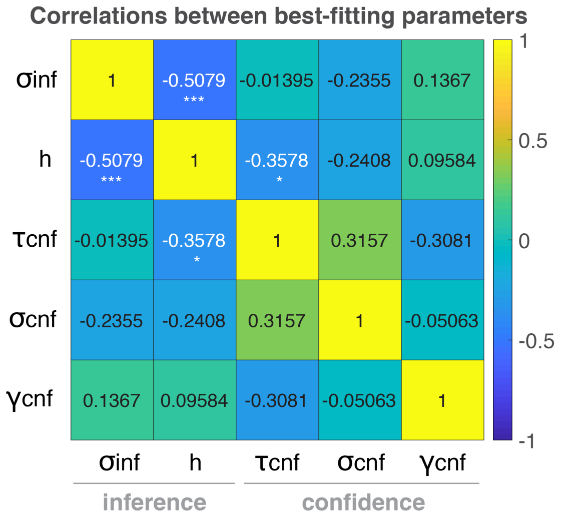


**Figure 4 supplement 2: Correlation between model parameters**. Individual best-fitting parameters are averaged across conditions. 1: inference noise; 2: hazard rate; 3: confidence threshold; 4: metacognitive noise; 5: confidence gain. ***$p$=.0025, *$p$=.041 indicates the significant correlations ($N$=33 participants). All other correlations were not significant (all abs(rho)<0.32, all $p$>.074).

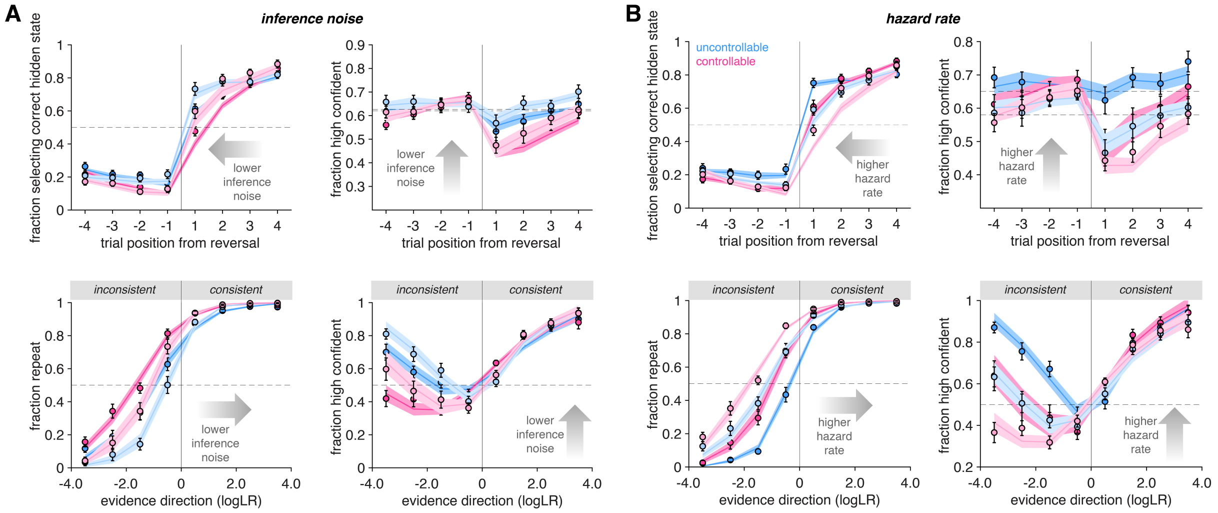


**Figure 4 supplement 3. Effect of inference noise and hazard rate parameters on choice and confidence patterns**

**A-B**) Effect of inference noise (**A**) and perceived hazard rate (**B**) parameters on choice and confidence patterns. Reversal and repetition curves for responses and confidence, with human data (circles) and simulations from the best-fitting computational model (shaded area) in uncontrollable (C-, blue) and controllable (C+, pink) conditions. Error bars represent SEM across participants (*N*=16 per group, median split of best-fitting inference noise and hazard rate respectively). Lighter colors indicate lower parameter values in all panels. Same conventions as Fig. 2 and Fig. 3.

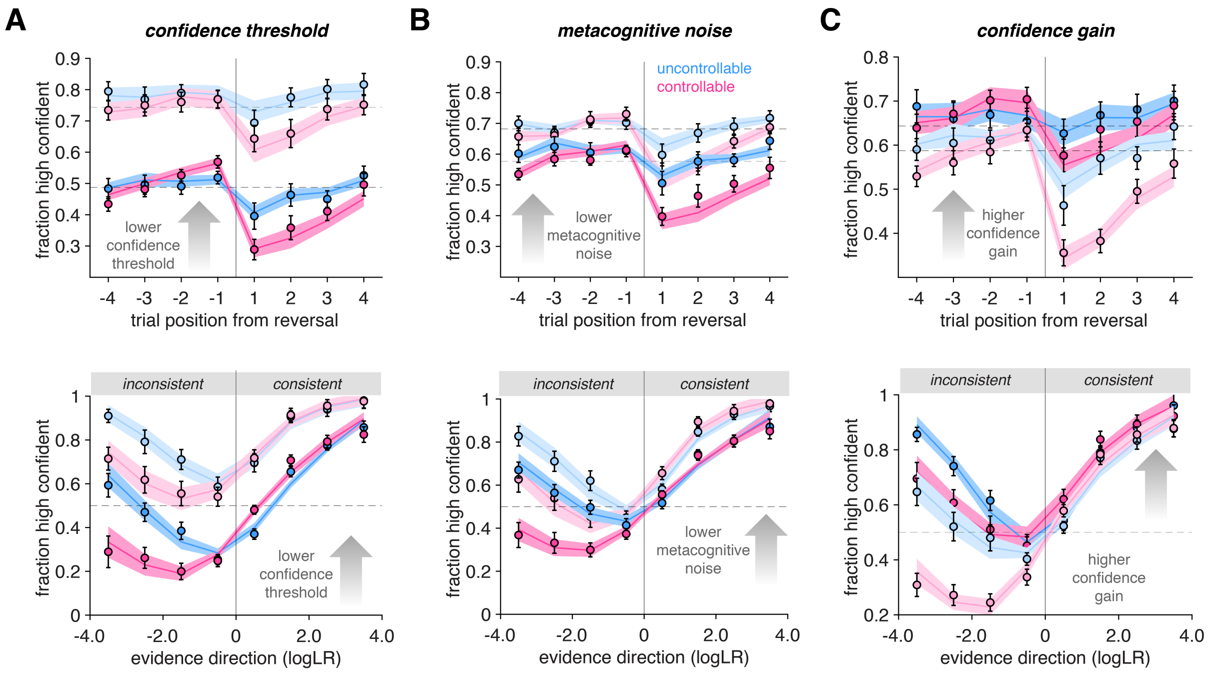


**Figure 4 supplement 4. Effect of confidence threshold, metacognitive noise, and confidence gain parameters on confidence patterns**

**A-C**) Effect of confidence threshold (**A**), metacognitive noise (**B**), and confidence gain (**C**) parameters on confidence patterns. Reversal and repetition curves for confidence, with human data (circles) and simulations from the best-fitting computational model (shaded area) in uncontrollable (C-, blue) and controllable (C+, pink) conditions. Error bars represent SEM across participants (*N*=16 per group, median split of best-fitting confidence threshold, metacognitive noise and confidence gain respectively). Note that choice curves are not shown because the three confidence parameters have no effect on choices, by definition. Lighter colors indicate lower parameter values in all panels. Same conventions as Fig. 2 and Fig. 3.

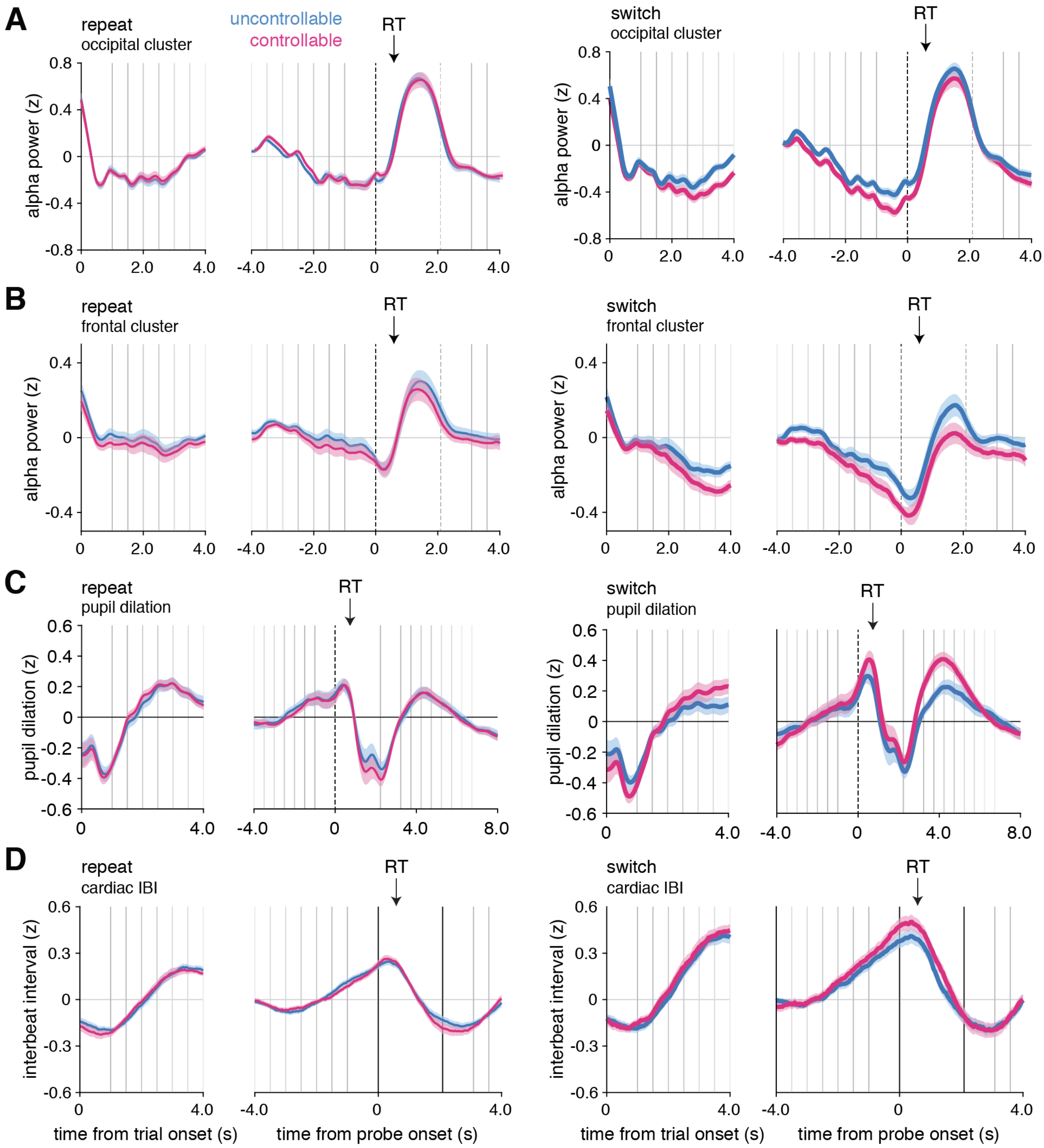


**Figure 6 supplement 1. Raw time-courses of MEG, pupil and IBI data**

**A**) Raw time-course of alpha-band power on repeat (left panels) and switch (right panels) trials in MEG for the occipital cluster. **B**) Raw time-course of alpha-band power on repeat (left panels) and switch (right panels) trials in alpha-band for the frontal cluster. **C**) Raw time-course of pupil dilation on repeat (left panels) and switch (right panels) trials. **D**) Raw time-course of cardiac interbeat interval on repeat (left panels) and switch (right panels) trials. In all panels, physiological responses were time-locked at the trial onset or the response probe onset. Vertical lines indicate trial events: trial onset, samples of the sequence, and response probe onset. Black arrows indicate the average response time across conditions and participants. Blue, uncontrollable (C-) condition; pink, controllable (C+) condition. *N*=24 participants from Experiment 4 for panels A-B-D. *N*=32 participants pooled across Experiments 1 and 2A for panel C.

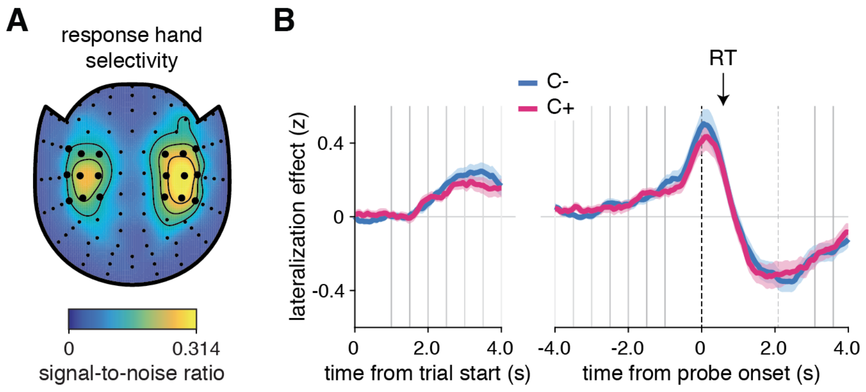


**Figure 6 supplement 2. Motor response preparation effects in alpha-band MEG (Experiment 4)**
**A**) Spatial topography of response hand selectivity at response onset, measured in units of signal-to-noise ratio. Large dots indicate the left and right channels used to compute a lateralization effect over time surrounding response probe onset. **B**) Contrast of switch minus repeat trials time-locked at the trial onset (left part) or the response probe onset (right part). Vertical lines indicate trial events: probe onset, samples of the sequence, and response probe onset. The black arrow indicates the average response time across conditions and participants (*N*=24). Blue, uncontrollable condition (C-); pink, controllable condition (C+).

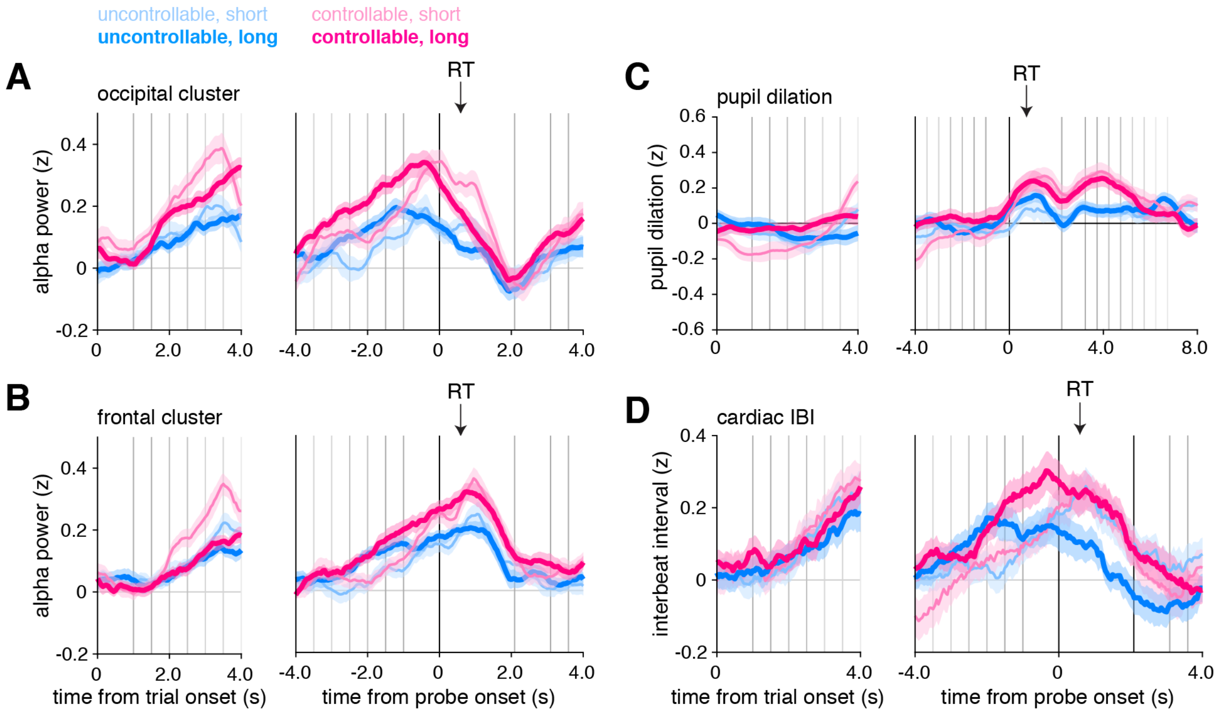


**Figure 6 supplement 3: Physiological analyses controlling for sequence length**

Contrast between repeat and switch trials for alpha-band power in the occipital cluster (**A**) and in the frontal cluster (**B**), for pupil dilation (**C**), and for cardiac interbeat interval (**D**) separately for short (2-4 samples) and long (6-8 samples) trial sequences. In all panels, physiological responses were time-locked to trial onset (left panels) or response probe onset (right panels). Lines represent averages and shaded areas SEM across participants. Vertical lines indicate trial events: probe onset, samples of the sequence and the response probe onset. Black arrows indicate average response time across conditions and participants. Blue: uncontrollable (C-) condition; pink: controllable (C+) condition. *N*=24 participants from Experiment 4 for panels A-B-D. *N*=32 participants pooled across Experiments 1 and 2A for panel C.

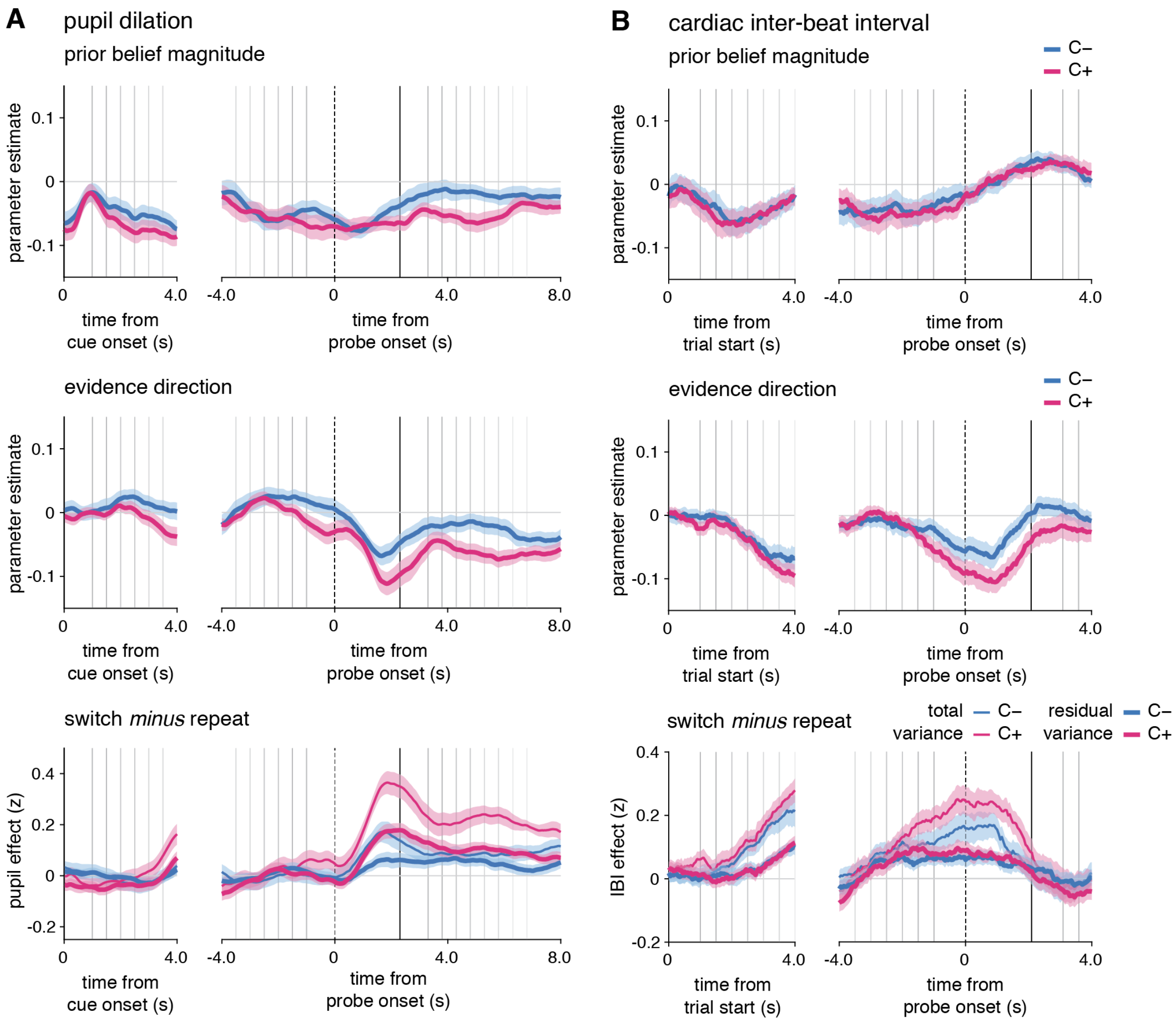


**Figure 8 supplement 1: Changes-of-mind effects after accounting for fluctuations in prior beliefs or evidence direction. A**) Effects of prior belief and evidence direction on pupil dilation and cardiac IBI. For pupil dilation (**A**) and cardiac IBI (**B**), contrast between repeat and switch trials time-locked at the trial onset (left panels) or at the response probe onset (right panels) after removing the variance due to fluctuations in prior belief and evidence direction (thick lines, residual variance) in the uncontrollable (blue, C-) and controllable (pink, C+) conditions. Raw contrasts are displayed for comparison (thin lines, total variance). Vertical lines indicate trial events: probe onset, samples of the sequence, and response probe onset (dotted line). The black arrow indicates the average response time across conditions and participants. total var.: total variance; residual var.: residual variance after accounting for the two model variables, prior belief and evidence direction.
